# Glucuronoxylomannan in the *Cryptococcus* species capsule as a target for CAR^+^ T-cell therapy

**DOI:** 10.1101/715045

**Authors:** Thiago Aparecido da Silva, Paul J. Hauser, Irfan Bandey, Tamara Laskowski, Qi Wang, Amer M. Najjar, Pappanaicken R. Kumaresan

## Abstract

The genus *Cryptococcus* comprises two major fungal species that cause clinical infections in humans: *C. gattii* and *C. neoformans*. To establish invasive human disease, inhaled Cryptococci must penetrate the lung tissue and reproduce. Each year, about 1 million cases of *Cryptococcus* infection are reported worldwide, and the infection’s mortality rate ranges from 20% to 70%. HIV^+^/AIDS patients are highly susceptible to *Cryptococcus* infection. Therefore, we hypothesized that CD8^+^ T cells could be redirected to target glucuronoxylomannan (GXM), a sugar present in the *Cryptococcus* species capsule, via expression of a GXM-specific chimeric antigen receptor (GXMR-CAR) for treatment of cryptococcosis. GXMR-CAR has an anti-GXM single-chain variable fragment followed by an IgG4 stalk, a CD28 transmembrane domain, and CD3-ς and CD28 signaling domains. After lentiviral transduction of human T cells with the GXMR-CAR construct, flow cytometry demonstrated that 82.4% of the cells expressed GXMR-CAR on their surface. To determine whether the GXMR-CAR^+^ T cells exhibited GXM-specific recognition, these cells were incubated with GXM for 24 h and examined using bright-field microscopy. Large clusters of proliferating GXMR-CAR^+^ T cells were observed, while no clusters were present in the control cells. Moreover, the interaction of GXM with GXMR-CAR^+^ T cells was detected via flow cytometry using a GXM-specific antibody. The ability of GXMR-CAR T cells to bind to the yeast form of *C. neoformans* was detected by fluorescent microscopy, but no binding was detected with NoDNA T cells. Furthermore, when GXMR-CAR^+^ T cells were administered to immunocompromised NSG mice infected with *C. neoformans* their *C. neoformans* burden was significantly lower than mock-transduced control T cell treated mice as shown via immunofluorescence using an anti-GXM antibody and Gomori methenamine-silver (GMS) staining of Titan cells in lung tissue. Thus, these findings demonstrated the effectiveness of GXMR-CAR^+^ T-cell therapy for cryptococcosis in a murine model.

**Author summary:** *Cryptococcus gattii* infects both immunocompetent and immunodeficient patients such as those with HIV/AIDS, while *C. neoformans* usually infects only immunocompromised patients. Every year, almost one million HIV/AIDS patients suffer from *Cryptococcus* fungal co-infection. At present, no curative treatment is available to treat cryptococcosis in chronic HIV/AIDS patients. The objective of this research was to develop novel “Bioengineered” *Cryptococcus* specific chimeric antigen receptor (CAR) CD8^+^ T-cells to target and kill *Cryptococcus*. By using a culture model, we demonstrated that the *Cryptococcus* specific CAR T cells were able to bind to the yeast form of *C. neoformans.* Using a mouse model of *Cryptococcus*, the *Cryptococcus* specific CAR treated group showed a significant reduction of fungal burden in lung tissue when compared to the control group. This gives new hope to HIV/AIDS patients suffering from cryptococcal infections.

## Introduction

*Cryptococcus gattii* and *Cryptococcus neoformans* are the two major *Cryptococcus* species that cause clinical infections in humans [1] due to the ability of basidiomycetous yeasts to grow at 37°C through asexual yeast budding in human and animal hosts. Upon inhalation of infectious propagules, *Cryptococcus* spp. penetrate the lung tissue and reproduce, causing pulmonary cryptococcosis, leading to the possibility of dissemination to other organs, most commonly the brain [2]. *C. gattii* infections occur in both immunocompetent and immunocompromised individuals, whereas *C. neoformans* infections are more common in immunosuppressed patients, such as those who have received organ transplants or have HIV infection/AIDs or hematological malignancies [3]. In mice infected with *C. neoformans*, CD4^+^ T-helper 1 (Th1) cells play an important role in fighting infection as demonstrated by the fact that interferon (IFN)-γ and interleukin (IL)-12–knockout mice had a higher mortality index than did mice wild-type for IFN-γ and IL-12 [4–7]. In addition, mice infected with *C. gattii* displayed reduced dendritic cell-mediated Th1/Th17 immune responses [8]. Although several fungal vaccine studies have demonstrated an essential role for CD8^+^ T cells in destroying host cells harboring intracellular fungus[9], not many studies have used CD8+ T cells for direct killing of extracellular fungal infection.

Our group was the first to explore the direct killing of extracellular fungi through the redirection of CD8^+^ T cells via expression of an engineered chimeric antigen receptor (CAR) to target a carbohydrate expressed on fungal cell walls [9, 10]. CARs are usually composed of four domains: 1) an extracellular antigen specific binding domain or domains, 2) a hinge or spacer region, 3) a transmembrane region, and 4) a cytoplasmic signaling region [11, 12]. Interest in using CAR^+^ T cells to treat diseases such as cancer has increased in recent years because CAR-based recognition enables T cells to identify proteins, glycoproteins, or glycolipids on the cell surface and release cytotoxic proteins which destroy the target cells in a manner that does not depend on the peptide-major histocompatibility complex expressed on antigen-presenting cells (APCs) [13]. CAR-dependent T-cell activation is achieved through the CAR endodomain, which is composed of CD28 or CD137 and CD3-ς [14, 15]. In most studies using CARs, researchers have administered CD19-targeted CARs for the treatment of B-cell malignancies. Recently, the U.S. Food and Drug Administration approved CAR^+^ T-cell therapies for leukemia and lymphoma [13, 16-20]. We demonstrated that T cells expressing a CAR containing the extracellular portion of Dectin-1 (D-CAR) targeted germinating fungal hyphae and inhibited hyphal growth of *Aspergillus* spp. [10]. However, the direct *Cryptococcus*-killing activity of CD8^+^ T cells has yet to be widely explored in the development of immunotherapy for infections with this pathogen.

Fungal cell wall glycans and exopolysaccharides are the first point of physical contact in fungal-host interactions, and these polysaccharides are common to multiple fungi [21]. Pattern recognition receptors present on innate immune cells recognize these glycans and eliminate fungal spores, germinating hyphae, and yeast. However, *Cryptococcus* spp. has exopolysaccharides that can mask these glycans that are recognized by immune cells. In the present study, we adapted a CAR to redirect engineered CD8^+^ T cells to target the polysaccharide capsule of *Cryptococcus* spp. The capsule of *Cryptococcus* spp. is a major virulence factor primarily composed of the sugar moiety glucuronoxylomannan (GXM), with lesser amounts of galactoxylomannan (GalXM) and mannoproteins [22–24]. One of the functions of the capsule is to provide protection from the recognition by the host immune system, which can be achieved by means such as preventing recognition and phagocytosis by the host innate immune cells, inhibiting the migration of phagocytes, and suppressing the proliferation of T cells [25]. Casadevall et al. [26] developed and characterized the murine monoclonal antibody (mAb) 18B7 that binds to *Cryptococcus* spp., promoting antibody dependent cellular toxicity mediated by innate immune cells such as macrophages. They found that administration of 18B7 to mice promoted clearance of the serum cryptococcal antigen GXM. Phase I clinical studies were performed using 18B7 to treat cryptococcosis in HIV^+^ patients. Titers of serum cryptococcal antigen decreased by a median of 2-3 fold in patients treated with of 1-2 mg/kg at 1-2 weeks post-infusion, and the treatment was well tolerated [26, 27].

The GXM-specific CAR (GXMR-CAR) was designed to achieve recognition of GXM using a single-chain variable fragment (scFv) originating from 18B7. We hypothesized that GXMR-CAR T cells would display cytotoxic activity against fungi expressing GXM on the cell wall. Our study demonstrates that engineered human T cells expressing GXMR-CAR bound to GXM in solution and interacted with heat killed *C. neoformans* yeast. The GXMR-CAR^+^ T cell treated NSG mice displayed significantly lower *C. neoformans* burden than mock-transduced control T cell treated mice.

## Results

### Construction of GXMR-CAR to target *Cryptococcus* spp

CAR^+^ T cells recognize antigens and undergo activation in a non-major histocompatibility complex-restricted manner, and investigators have successfully used them to trigger immune response against cancer cells. We adopted this approach to generate CAR^+^ T cells that target *Cryptococcus* spp., which cause infections in immunocompromised (*C. neoformans*) and immunocompetent (*C. gattii*) individuals. GXM, is the major virulence factor in the *Cryptococcus* spp. capsule and protects the yeast from the host immune cell attack. The light and heavy chains were selected from the murine monoclonal antibody 18B7 [26] to generate a scFv that specifically recognizes GXM present in the *Cryptococcus* spp. capsule.

To generate the GXMR-CAR construct, the DNA sequence of the GXM-specific scFv (scFv-18B7) was fused to modified human IgG4 hinge and Fc regions, transmembrane and co-stimulatory domains of CD28, and the signaling domain of CD3-ς (Fig 1A) as reported previously [10]. HEK-293FT cells were used to make GXMR-CAR^+^ viral particles by transfecting them with three plasmids: pMD2.G for the envelope, psPAX2 for packaging, and GXMR-CAR, according to the manufacturer’s instructions as described in the Materials and Methods section. The success of transfection was determined by analyzing green fluorescent protein (GFP) expression after 24 h (Fig 1B) at 100× and 200× magnification.

**Fig 1.**
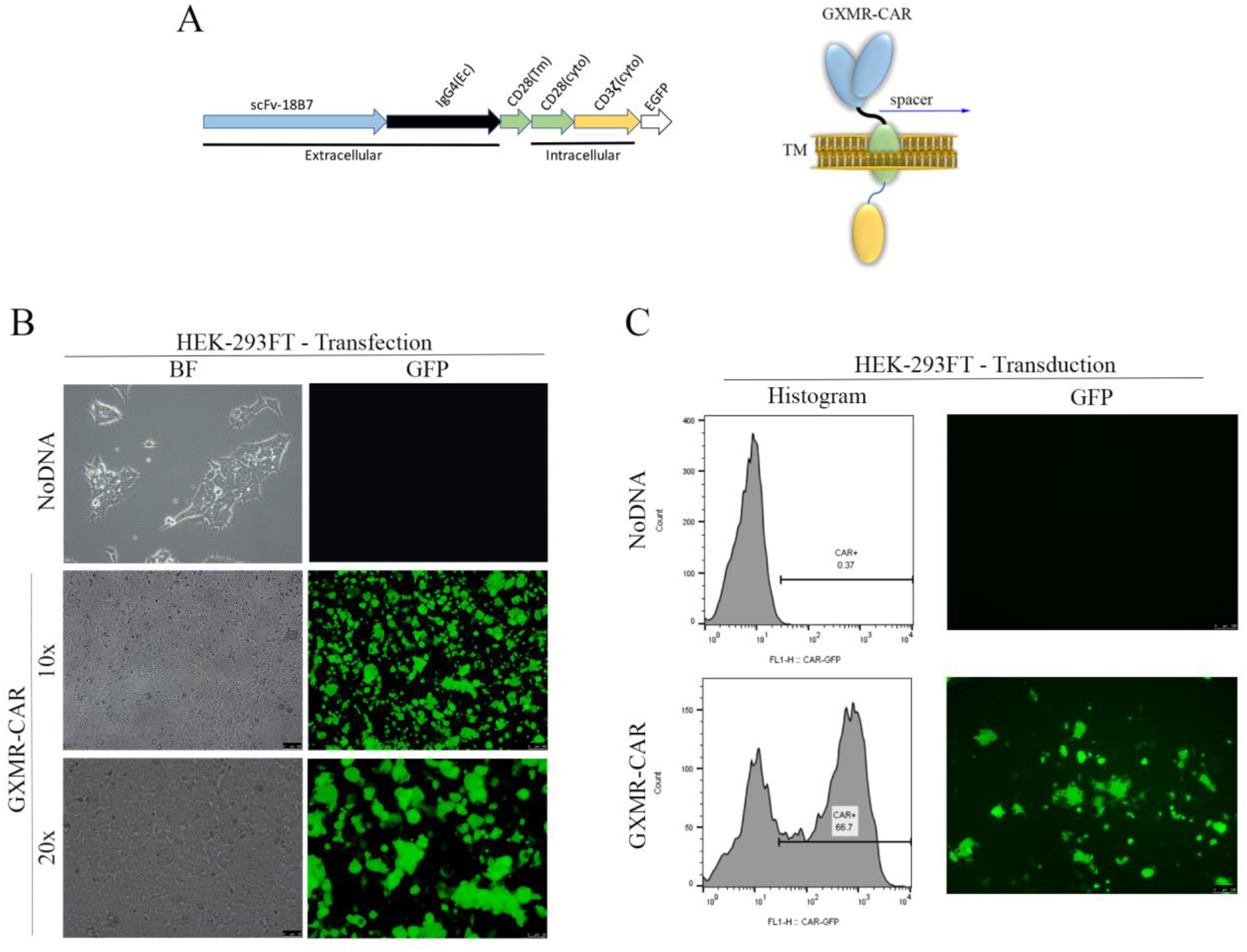
Construction of a GXMR-CAR that targets *Cryptococcus* spp. (**A**) The DNA sequence (left) and schematic representation (right) of the GXMR-CAR targeting *Cryptococcus* spp. has a scFv portion derived from the anti-GXM mAb 18B7. The DNA is composed of four domains: the signaling domains of CD3-ς and CD28 (yellow and green regions, respectively), the transmembrane (Tm) domain of CD28 (green region), the constant region of IgG4 (black line), a scFv portion from 18B7 (blue region), and an enhanced GFP (EGFP) portion for detection via fluorescence (white region). The DNA sequence was subcloned into a lentiviral vector backbone. The LV-GXMR-CAR construct was used for transfection and transduction. Ec, extracellular binding domain; cyto, cytoplasmic signaling region. (**B**) HEK-293FT cells (1 × 10^6^ cells/ml) were used to make GXMR-CAR^+^ viral particles via transfection with all three plasmids combined: LV-GXMR-CAR, pMD2.G, and psPAX2, using Lipofectamine. Mock-transduced HEK-293FT cells (NoDNA) were used as negative controls. HEK-293FT cells were visualized using fluorescence microscopy in bright-field (BF) and green (GFP) channels at 100× and 200× magnification 24 h after transfection. (**C**) GXMR-CAR^+^ viral particles were used to transduce HEK-293FT cells. HEK-293FT cells were evaluated via flow cytometry and expressed in a histogram (left). HEK-293FT cells were visualized using fluorescence microscopy in brightfield and GFP channels at 100× magnification (right) 3 days after transduction. Mock-transduced negative control cells (NoDNA) did not receive the GXMR-CAR construct.

After the viral particles containing GXMR-CAR were generated, HEK-293FT cells were infected with the particles to evaluate the GXMR-CAR transduction efficiency. After three days in culture, the GFP positive cells were identified by using fluorescence microscopy (Fig 1C, right). and the GFP expression in transduced cells was quantified using flow cytometry (Fig 1C, left) A high efficiency of transduction of HEK-293FT cells with GXMR-CAR^+^ viral particles was observed, as 67% of the cells were GFP^+^ (Fig 1C).

### GXMR-CAR expression on human T cells

After GXMR-CAR was successfully expressed in HEK-293FT cells, the same protocol was utilized for the transduction of human T cells. Initially, peripheral blood mononuclear cells (PBMCs) were stimulated with an antibody cocktail (anti-CD3/CD28) combined with IL-2 for 3 days. Activated human T cells were transduced with GXMR-CAR^+^ viral particles and after 3 days post-transduction, GXMR-CAR^+^ T cells were enriched by sorting based on GFP expression, and expansion of GXMR-CAR^+^ T cells was performed as described in Materials and Methods. After 3 days of expansion, 82.4% of the human GXMR-CAR^+^ T cells expressed GFP as determined using flow cytometry (Fig 2A). To demonstrate expression of GXMR-CAR on the surface of T cells, an anti-human Fc antibody (Fc-γ fragment-specific) was used to bind to the spacer region of IgG4 (as shown in Fig 1A) localized on their surface. The results show that 77.8% of the GXMR-CAR^+^ T cells were positive for the antibody (Fig 2B), confirming the expression of GXMR-CAR on the T-cell surface. Expanded GXMR-CAR^+^ T cells were evaluated by flow cytometry to determine if the stem cell memory, central memory, and effector memory T-cell subsets, which are critical for long-term immune memory, were prevalent in GXMR-CAR^+^ T cells (Fig 2 C, D).

**Fig 2.**
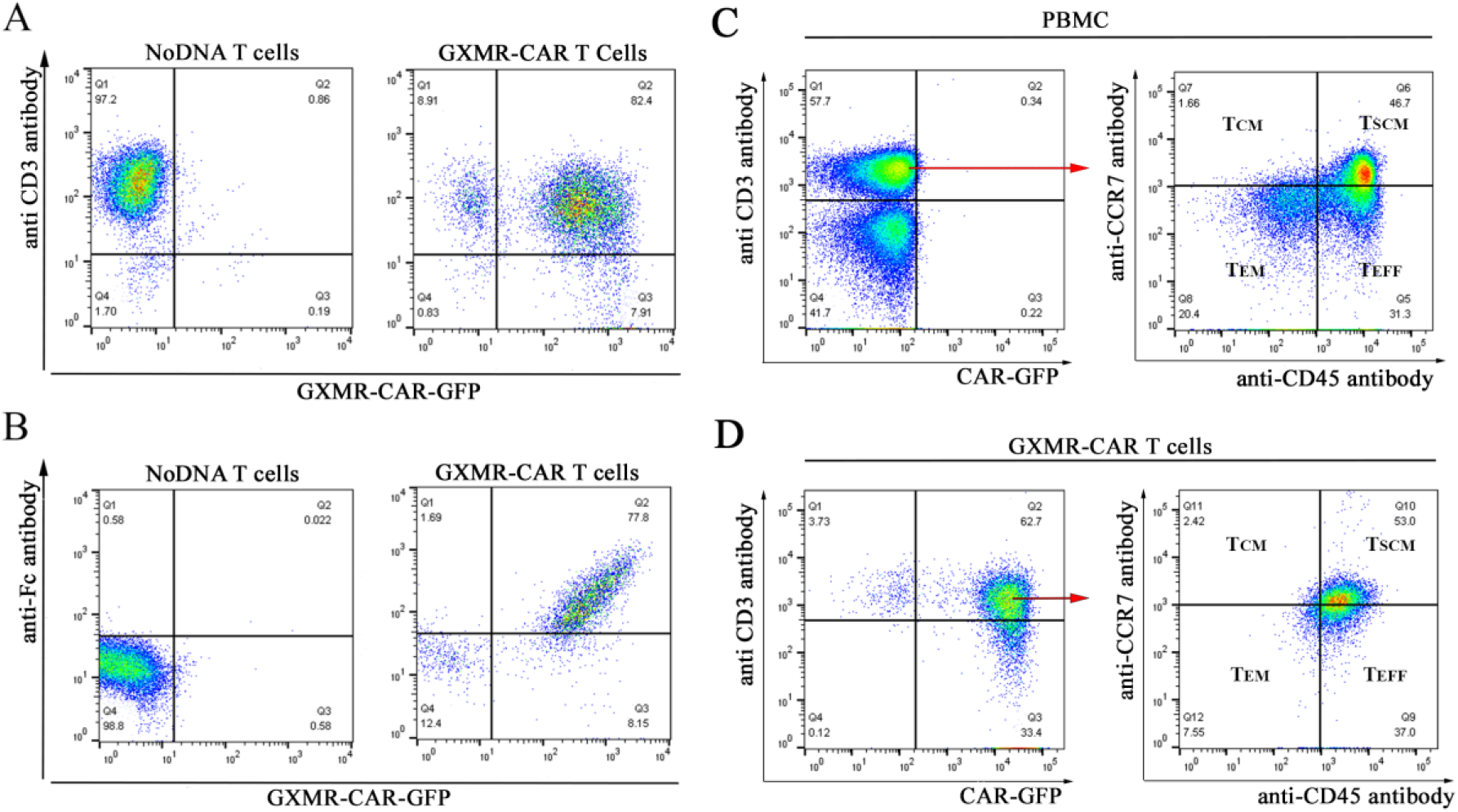
Expression of GXMR-CAR on the surface of human T cells. (**A**) Flow cytometry plots showing anti-CD3 antibody-positive T cells on the y-axis and GXMR-CAR-GFP– expressing T cells on the x-axis. The cells double-positive for the antibody and GFP (82.4%) are GXMR-CAR^+^ GFP-expressing T cells. (**B**) Flow cytometry plots showing T cells positive for an anti-human Fc antibody on the y-axis and GXMR-CAR-GFP–expressing cells on the x-axis. The cells double-positive for the antibody and CAR-GFP (77.8%) were considered GXMR-CAR^+^ T cells. Mock-transduced negative control T cells (NoDNA) did not receive the GXMR-CAR construct. **Phenotypic analysis of GXMR-CAR^+^ T cells (C, D):** This analysis of GXMR-CAR^+^ T cells stimulated using an ImmunoCult human CD3/CD28 T-cell activator was performed after 30 days of *in vitro* expansion of the cells. (**C**) PBMCs and (**D**) GXMR-CAR^+^ T cells were stained with a mixture of anti-CD3-PE, anti-CD8-PerCP, anti-CCR7-APC, and anti-CD45RA-PE-Vio770 antibodies, and CAR^+^ T cells were evaluated according to GFP expression. CD3^+^CD8^+^ T cells were gated to evaluate the memory cell subsets using anti-CCR7 and anti-CD45RA antibodies. The memory cell subsets were classified as follows: T_CM_, central memory T cells; T_EM_, effector memory T cells; T_EFF_, effector T cells; T_SCM_, stem cell memory T cells.

### GXMR-CAR^+^ T cells interact with soluble GXM from the *Cryptococcus spp.* capsule

To determine whether GXMR-CAR^+^ T cells recognize GXM polysaccharide from *Cryptococcus* spp., GXMR-CAR^+^ T cells were incubated with a preparation of polysaccharide antigens from the capsule of *Cryptococcus* spp. After 24h of incubation with polysaccharide antigens from *Cryptococcus spp.*, cell clusters were observed by bright-field microscopy in wells treated with GXMR-CAR T cells, while no cell clusters were detected upon incubation of NoDNA T cells (mock transduced cells) with the polysaccharide antigen (Fig 3A). Similar types of cell clusters were observed in the anti-CD3/CD28 antibody treated positive control, while no cell clusters were observed in the medium alone negative control wells (Fig 3A). These results suggest that GXMR-CAR^+^ T cells interact with polysaccharide antigens in the capsule of *Cryptococcus* spp., promoting the clustering of GXMR-CAR^+^ T cells.

**Fig 3.**
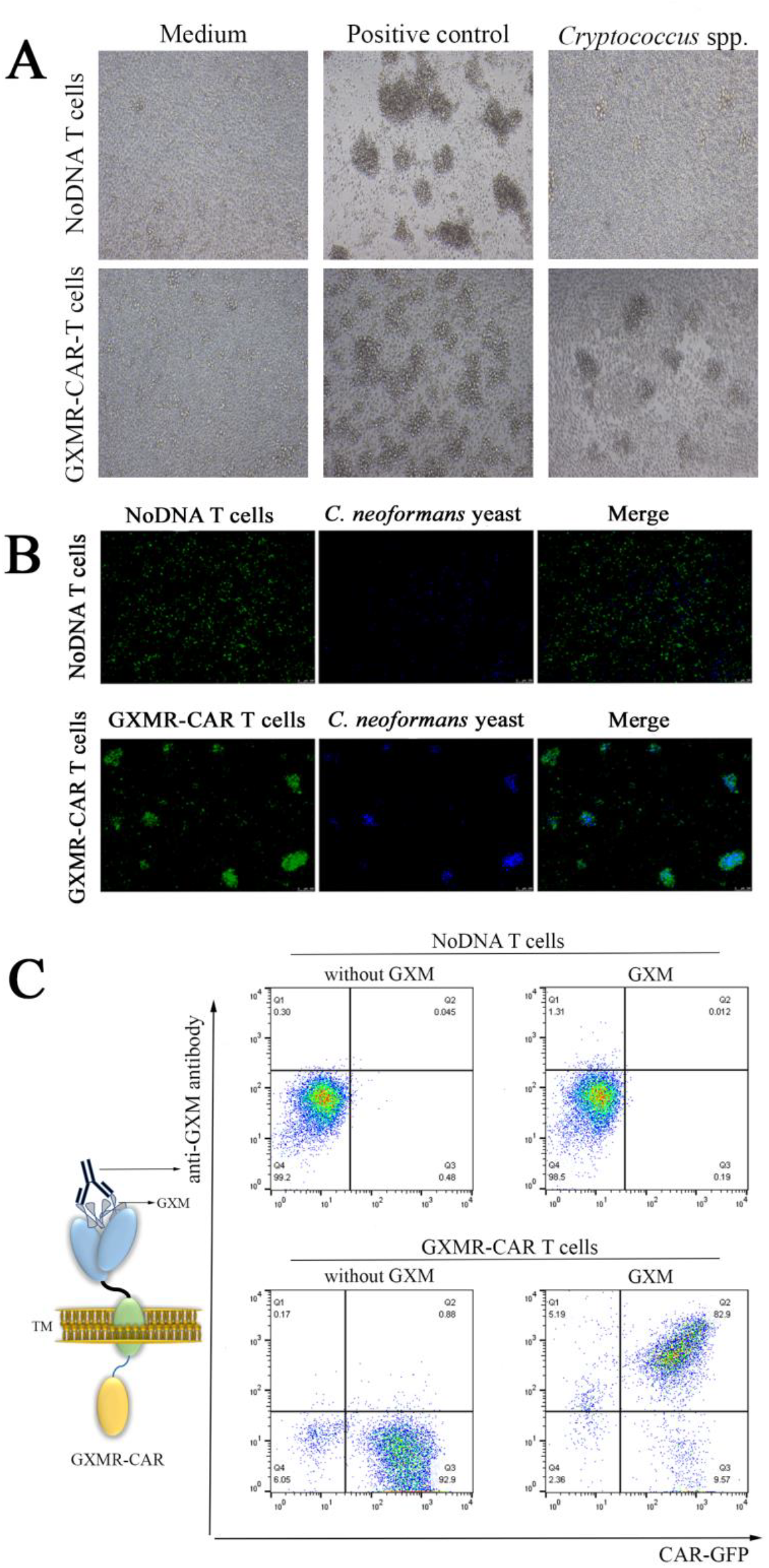
Recognition of GXM from *Cryptococcus* spp. by GXMR-CAR^+^ T cells. (**A**) GXMR-CAR^+^ and NoDNA T cells were incubated with a preparation of polysaccharide antigens from the capsule of *Cryptococcus* spp. at a 1:100 dilution. After 24h, the images were acquired by bright-field microscopy at 400x magnification. Positive control T cells were activated using an ImmunoCult human CD3/CD28 T-cell activator (STEMCELL Technologies) following the manufacturer’s instructions. T cells incubated with medium alone were used as negative control cells. (**B**) GXMR-CAR^+^ T cells and NoDNA T cells (mock-transduced, negative controls) were labeled with carboxyfluorescein succinimidyl ester and incubated with *C. neoformans* yeast labeled with Calcofluor-white. The interaction of GXMR-CAR^+^ T cells (green) and NoDNA T cells (green) with *C. neoformans* yeast (blue) was evaluated using fluorescence microscopy. The bottom right image shows the cell clusters formed by co-localization of GXMR-CAR^+^ T cells and *C. neoformans* yeast. NoDNA T cells (top right image) did not display co-localization. (**C**) GXMR-CAR^+^ and NoDNA T cells were assayed with polysaccharide antigens from the capsule of *Cryptococcus* spp., and the interaction of GXM with the cell surface was evaluated via flow cytometry using an anti-GXM mAb 18B7 clone stained with a phycoerythrin-conjugated secondary antibody. The cells double-positive for 18B7 and GXMR-CAR-GFP demonstrated the interaction between GXMR-CAR^+^ T cells and GXM from *Cryptococcus* spp. The NoDNA T cells and GXMR-CAR^+^ T cells not incubated with polysaccharide antigens from the capsule of *Cryptococcus* spp. (without GXM) were used to demonstrate the absence of nonspecific binding of 18B7 to cells.

The recognition of GXM by GXMR-CAR^+^ T cells allowed to investigate whether GXMR-CAR^+^ T cells target *C. neoformans* yeast. The *C. neoformans* yeast was stained with Calcofluor-white (blue), and incubated with GXMR-CAR^+^ T cells (green) or NoDNA T cells (negative control; green), and their interaction was visualized using fluorescence microscopy. The results show that GXMR-CAR^+^ T cells co-localized with *C. neoformans* yeast (Fig 3B, right), while the NoDNA T cells did not interact with the yeast (Fig 3B, left). Moreover, in GXMR-CAR^+^ T cell-treated wells, *C. neoformans* yeast was trapped inside T-cell clusters, whereas in wells with NoDNA T cells, the cells did not interact with the yeast, and no cell clusters were observed. These results demonstrate that GXMR-CAR^+^ T cells can recognize *Cryptococcus* spp. yeast and form complexes with it. To corroborate these findings, additional experiments were conducted to determine the specificity of GXMR-CAR^+^ T cells for GXM. Polysaccharide antigens from the capsule of *Cryptococcus* spp. were incubated with GXMR-CAR^+^ T cells, and an anti-GXM mAb 18B7 was added to detect GXM on the surface of GXMR-CAR^+^ T cells using flow cytometry (Fig 3C, right). The recognition of soluble GXM by GXMR-CAR^+^ T cells was demonstrated by cells that were double-positive for CAR-GFP and the anti-GXM mAb 18B7 clone (Fig 3C), whereas NoDNA T cells did not recognize GXM. These findings confirmed that the GXMR-CAR–expressing T cells targeted GXM, a major compound in the *Cryptococcus* spp. capsule.

### Targeting efficacy of GXMR-CAR^+^ T cells in a pulmonary *Cryptococcus* infection model

NOD *Scid* gamma (NSG) mice were used for these studies because they are deficient in mature B, T, and natural killer cells. Therefore, they are unable to target and destroy infused GXMR-CAR^+^ T cells. Pulmonary fungal infection was established in the mice by infecting their lungs with *C. neoformans* yeast (1 × 10^5^/mouse) via intranasal infusion [28]. The mice were placed into 3 groups according to the infusions they received: GXMR-CAR^+^ T cells, NoDNA T cells, and phosphate-buffered saline (PBS). The mice were infused with 5 million GXMR-CAR^+^ T cells, NoDNA T cells, or PBS alone via tail vein injection at 1 and 4 days after infection. Eight days after infection, the mice were humanely euthanized and entire sections of lung tissue were evaluated by immunofluorescence using an anti-GXM antibody and imaging was performed at 400x (Fig. 4A). More than 2000 images were analyzed using the automated inForm^®^ Cell Analysis^™^ software to quantify the area of GXM-FITC stained Titan cells (Fig. 4B, red shaded region) and total area of lung tissue (green shaded region) on each slide. A smaller area of GXM positive Titan cells (hallmark of *Cryptococcus* infection in the lung) was observed in the GXMR-CAR samples (0.021 mm^2^ GXM, 57.55 mm^2^ lung) compared to the negative controls (PBS: 0.37 mm^2^ GXM, 39.2 mm^2^ lung; NoDNA: 0.066 mm^2^ GXM, 60.0 mm^2^ lung), indicating a reduced fungal burden due to treatment with GXMR-CAR^+^ T cells (Fig 4). These results were supported by Gomori methenamine-silver (GMS) staining of Titan cells in lung tissue sections obtained from the mice in the PBS, GXMR-CAR^+^ T-cell, and NoDNA T-cell groups. A lower fungal burden was observed in the GXMR-CAR^+^ T-cell group compared with the PBS group (Fig 4C). In addition, ImageJ was used to analyze the nine anti-GXM immunofluorescence images shown in Fig 5A, and determine the area of GXM positive staining in the lung tissues. Figure S1 shows some of the processing steps involved in processing the images using ImageJ. The ImageJ analysis demonstrated a significant reduction in the area of GXM staining in the GXMR-CAR T cell treated group compared to the PBS control. Corroborating our *in vitro* results, these data demonstrate that intravenously infused GXMR-CAR^+^ T cells directly target *C. neoformans* and control its spread in the lungs of infected mice.

**Fig 4.**
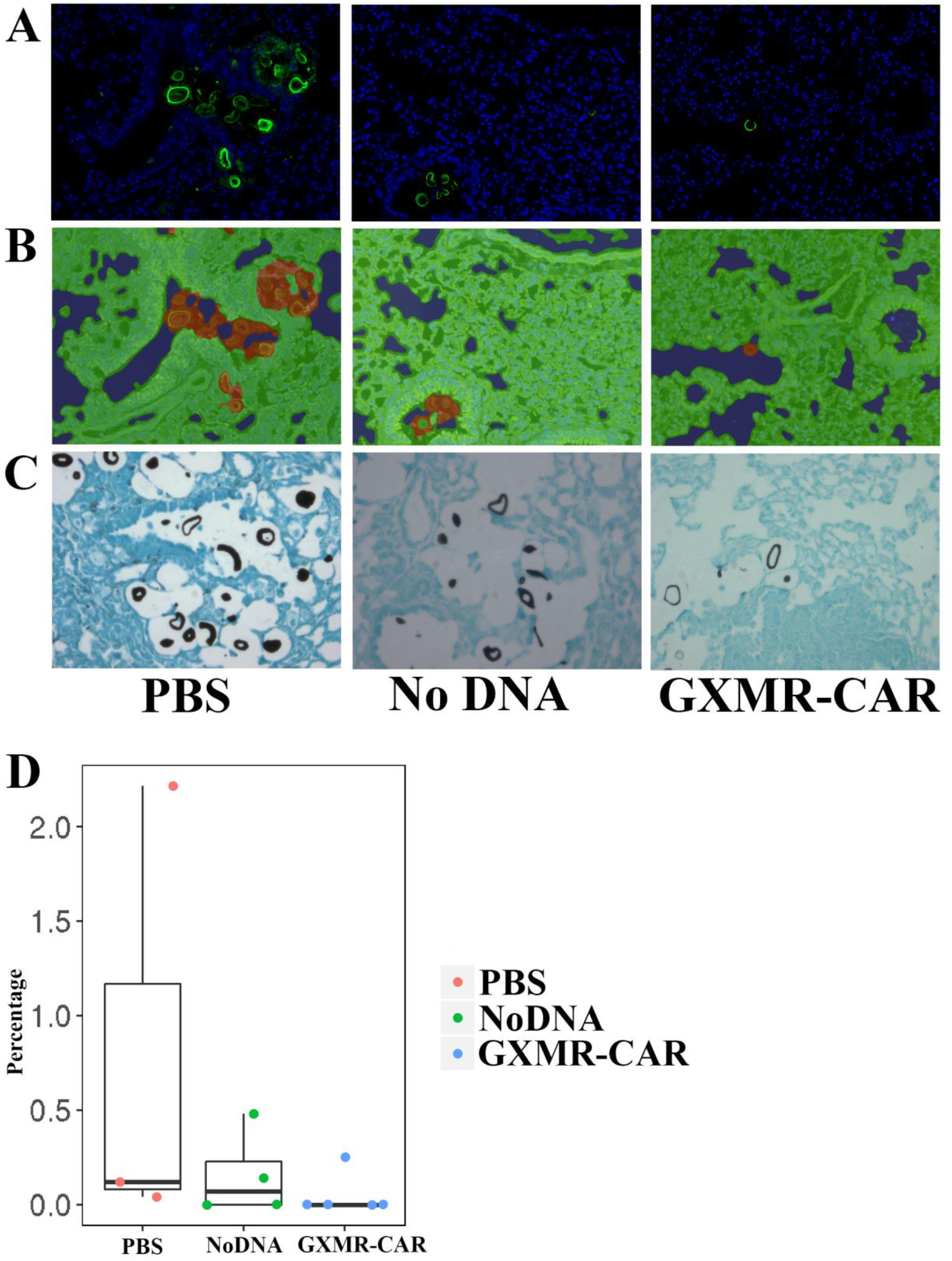
The automated inForm^®^ Cell Analysis^™^ software shows reduced area of GXM positive Titan cells in in the lungs of *C. neoformans* infected mice treated with GXMR-CAR^+^ T cells. The lungs of NSG mice (5 per group) were infected with *C. neoformans* yeast (1 × 10^5^/mouse) via intranasal infusion. Mice were infused intravenously with 5 million GXMR-CAR^+^ T cells, NoDNA T cells or PBS alone on days 1 and 4 after infection. Eight days after infection, the mice were humanely euthanized. (**A**) Paraffin-embedded lung tissue sections from the mice were incubated with an anti-GXM antibody and anti-mouse-FITC secondary to detect Titan cells as described in Materials and Methods. Images were acquired at 400× magnification using a Vectra Polaris microscope. Spectrally unmixed images showing DAPI labeled nuclei (Blue) and GXM-FITC labeled Titan cells (green) are shown. (**B**) The automated inForm^®^ Cell Analysis^™^ software was utilized to quantify the area of GXM positive staining and the total area of lung tissue. Images show GXM positive Titan cells (red), lung tissue (green), non-tissue (blue). (**C**) Paraffin-embedded lung tissue sections from the mice were stained with Gomori methenamine-silver (GMS) to analyze Titan cells (black circles) as described in Materials and Methods. The images were acquired at 200× magnification using an Olympus CKX41 microscope. (D) Graph displaying results of GXM immunofluorescence assay analyzed by the automated inForm^®^ Cell Analysis software. The percentages of the “area of GXM positive staining / total area of lung tissue” for each mouse per group were compared using ANOVA. Pair-wised comparisons were conducted using Tukey’s HSD (honestly significant difference) test. Statistical significance was defined as P value < 0.05.

**Fig 5.**
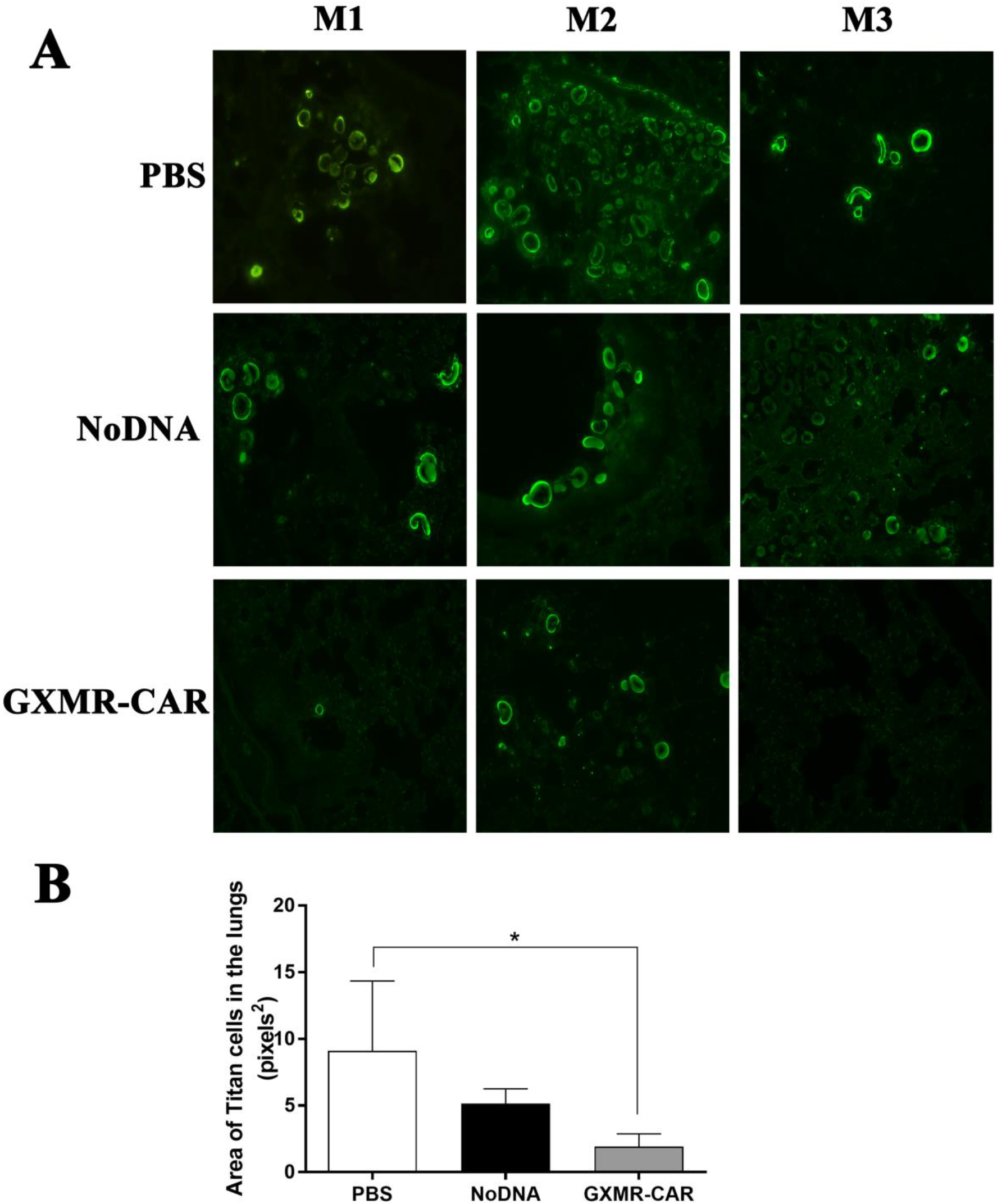
ImageJ analysis of anti-GXM immunofluorescence assay shows reduced area of GXM positive Titan cells in the lungs of *C. neoformans* infected mice treated with GXMR-CAR^+^ T cells. The lungs of NSG mice (5 per group) were infected with *C. neoformans* yeast (1 × 10^5^/mouse) via intranasal infusion. Mice were infused intravenously with 5 million GXMR-CAR^+^ T cells, NoDNA T cells or PBS alone on days 1 and 4 after infection. Eight days after infection, the mice were humanely euthanized. (**A**) Paraffin-embedded lung tissue sections from the mice were incubated with an anti-GXM antibody and anti-mouse-FITC secondary to detect Titan cells as described in Materials and Methods. Images were acquired at 200× magnification using an Olympus CKX41 microscope. GXM-FITC labeled Titan cells (green) are shown. (**B**) ImageJ software was used to quantify the area of GXM positive staining in lungs from the nine the images that are shown in Fig 5A. A statistically significant reduction in the area of GXM stained Titan cells was demonstrated in GXMR-CAR T cell treated mice compared with PBS controls.

**Fig 6.**
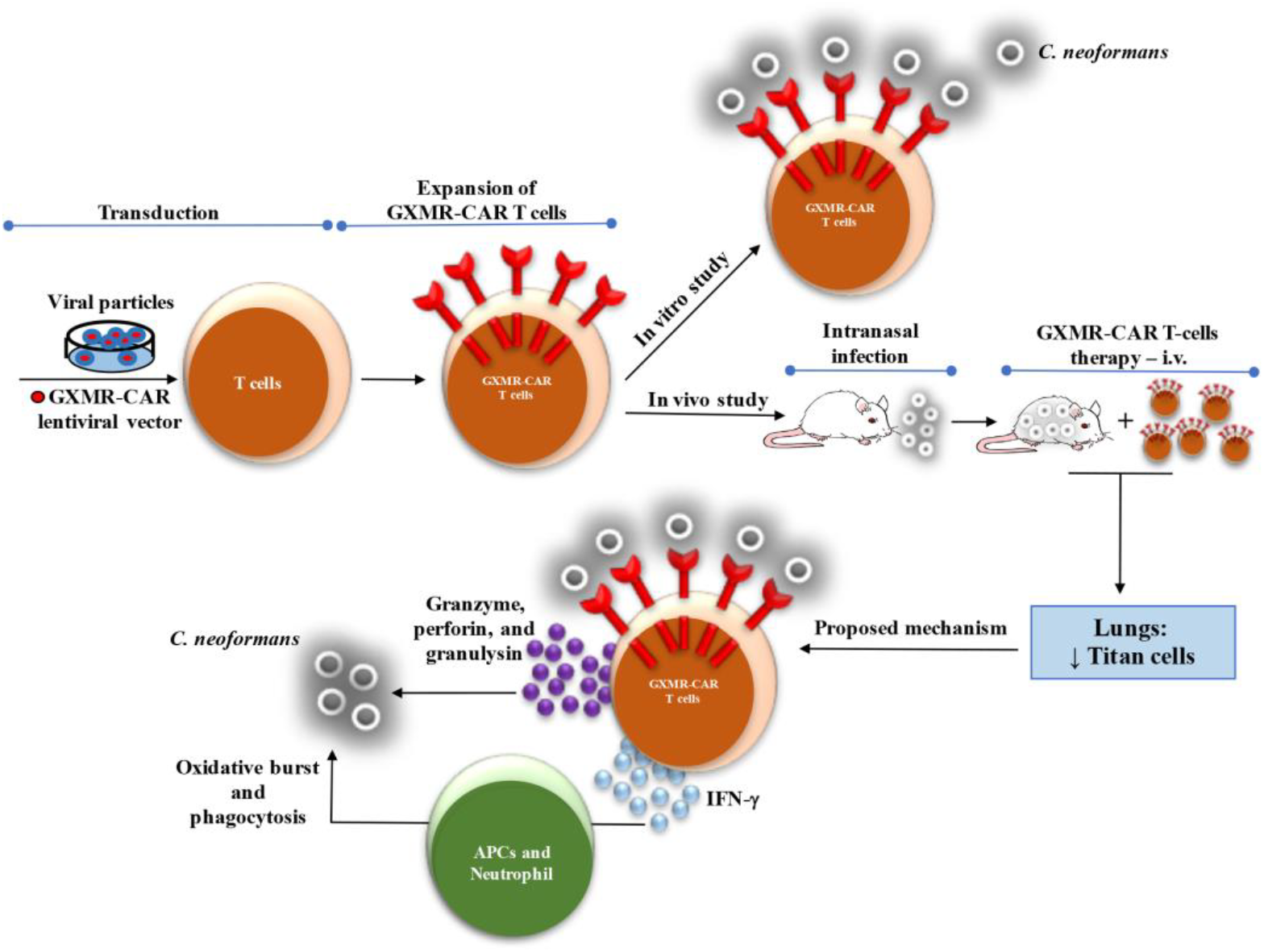
Approach to targeting *C*. *neoformans* with CAR^+^ T cells. Schematic showing that T cells engineered to express GXMR-CAR via lentiviral transduction can target *C. neoformans*. *In vitro* and *in vivo* studies demonstrate that GXMR-CAR^+^ T cells interact with *C. neoformans* yeast and that GXMR-CAR^+^ T cells infused in NSG mice infected with *C. neoformans* promoted a reduction in the number of Titan cells in the lungs. We suggest that GXMR-CAR^+^ T cells use two major mechanisms to control cryptococcosis: 1) activation of neutrophils and APCs by proinflammatory mediators released by GXMR-CAR^+^ T cells and 2) production of granzyme, perforin, and granulysin by GXMR-CAR^+^ T cells acting on the *C. neoformans*.

## Discussion

Physicians have successfully used CAR^+^ T-cell therapy to treat hematological malignancies, with an overall success rate greater than 80% [29]. In 2017, the U.S. Food and Drug Administration approved two CD19-targeting CARs (tisagenlecleucel [Kymriah] and axicabtagene ciloleucel [Yescarta]) for treatment of chronic lymphoblastic leukemia [19], pediatric relapsed or refractory acute lymphoblastic leukemia, and relapsed or refractory large B-cell lymphoma [29]. The redirection of T cells by a CAR targeting a carbohydrate expressed on fungal cell walls was first reported by Kumaresan et al. [10]. In the present study, we are the first group to develop CAR^+^ T cells targeting the yeast virulence factor GXM, a carbohydrate present in the *Cryptococcus* spp. capsule. The extracellular domain of GXMR-CAR^+^ T cells was derived from the murine mAb 18B7, which binds to GXM and has a high affinity for *Cryptococcus* spp. In a phase I dose escalation trial, researchers tested treatment of cryptococcal meningitis in HIV^+^ patients with 18B7 [26, 27]. They observed transient decreases in serum cryptococcal antigen levels in patients receiving 18B7 doses of 1-2 mg/kg and that doses up to 1 mg/kg were well tolerated. However, the use of CAR T cell therapy has not been explored for treating cryptococcosis. We designed GXMR-CAR as a second-generation CAR with a similar structure to CD19RCAR, which has undergone in clinical trials (www.clinicaltrials.gov, NCT01318317 and NCT01815749), except that the scFv domain, which targets GXM, was swapped with the scFv that targets CD19. The lentiviral vector that was used for genomic insertion of CAR constructs was selected because (1) it has been used in clinical trials, (2) large quantities of CAR T cells can be produced within 7-10 days of transduction, and (3) more memory cells are produced (T_CM_ and T_EM_ subtypes) when compared to non-viral methods of CAR T-cell production. GXMR-CAR T cells were able to bind to GXM after incubation with *C. neoformans* polysaccharide antigens, and recognize *C. neoformans* yeast in both *in vitro* and *in vivo* studies. These findings demonstrate that the scFv domain of the GXMR-CAR^+^ T-cells retains its targeting potential against *C. neoformans* and could be used to control cryptococcosis.

The currently available antifungal drugs used to combat cryptococcosis consist of a combination of amphotericin B deoxycholate or liposomal amphotericin B and 5-fluorocytosine administered for 2 weeks. Fluconazole is recommended for use during the consolidation and maintenance phases, but this antifungal drug requires a long treatment course, which reduces patient compliance and/or tolerability [30, 31]. The treatments of cryptococcosis and other invasive fungal infections should display rapid fungicidal activity and completely eliminate the fungus from the host system. The rising number of antibiotic resistance strains of fungi, such as *C. gattii* [32] and *Candida auris* [33] also poses a significant threat to health. From 2004-2007 there were 83 cases of *C. gattii* infection in Washington and Oregon, with a mortality rate of 33%. It is thought that antibiotic resistance may have contributed to the poor response rate of antibiotic therapy [32]. To tackle such circumstances other treatment options such as fungal immunotherapy are needed to treat these deadly diseases.

To improve the currently available treatment options for cryptococcosis, combining immunomodulators with existing antifungal drugs may be beneficial. Along these lines, researchers evaluated the effect of administration of adjuvant recombinant IFN-γ in 75 HIV^+^ patients with acute cryptococcal meningitis, which led to more rapid sterilization of cerebrospinal fluid in 36% of the treated group compared with 13% of the placebo controls. Moreover, the addition of recombinant IFN-γ to amphotericin B in HIV^+^ patients with cryptococcal meningitis caused a significant reduction of *Cryptococcus* spp. in cerebrospinal fluid [34]. Recently, a research group used a vaccination strategy to control cryptococcosis using glucan particles as a delivery system to carry protective protein antigens from *C. neoformans* or *C. gattii*. This approach induced CD4^+^ T-cell responses in the lungs of vaccinated mice, providing partial protection after challenge with *C. neoformans* or *C. gattii* [35]. These studies demonstrated that polarization of adaptive immune cells is critical to an appropriate immune response against cryptococcosis.

Another immunomodulatory treatment strategy is adoptive T-cell therapy, which can induce cell-mediated immunity against invasive fungal infections. CD4^+^ T cells differentiate into Th1 and Th17 cells that mainly secrete IFN-γ and IL-17, respectively, activating innate immune cells to fight fungal infections [36, 37]. In addition, the proinflammatory cytokines produced by Th1 and Th17 cells activate B cells, resulting in the secretion of antigen-specific antibodies against fungi. Moreover, CD8^+^ T cells differentiate into Tc1 and Tc17 cells, which produce IFN-γ and IL-17, contributing to the recruitment of innate immune cells involved in antifungal defense. The major mechanism that CD8^+^ T cells use to control fungal infections is the targeted release of cytotoxic factors that act directly on fungi, such as perforin, granulysin, and granzyme [9]. The antifungal activity of adoptive T-cell therapy can be improved by redirecting T-cell specificity using a CAR that recognizes specific fungal antigens. In this context, Kumaresan et al. [10] demonstrated that engineered CAR^+^ T cells containing an extracellular domain of the carbohydrate recognition domain of Dectin-1 (D-CAR–expressing T cells) were capable of targeting β-glucan–expressing fungi. *In vitro* and *in vivo* studies showed that D-CAR–expressing T cells target germinating *Aspergillus* hyphae and inhibit aspergillosis dissemination by using the cytolytic machinery of the genetically modified T cells [10].

These findings led us to develop CAR^+^ T cells expressing an scFv from the mAb 18B7 to target *Cryptococcus* spp. that contain a polysaccharide capsule predominantly comprised of GXM [38]. Because the compounds in the *Cryptococcus* spp. capsule impair the recognition of pathogen-associated molecular patterns by pattern recognition receptors, the β-glucans, mannoproteins, and chitin are poorly recognized by the immune system. GXMR-CAR expressed on human T cells recognizes the soluble form of GXM and targets *C. neoformans* yeast, demonstrating the specificity of GXMR-CAR to this pathogen. The major fungi responsible for invasive fungal infections, *Candida* and *Aspergillus* species, lack GXM and have different polysaccharide compositions in the outer cell wall [21], while GXMR-CAR specifically targets only fungi that have GXM in the outer cell wall. Our *in vivo* studies demonstrated that GXMR-CAR^+^ T cells efficiently reduced the number of Titan cells in the lungs of mice infected with *C. neoformans* when compared to the negative control group, as demonstrated by immunofluorescence and histochemistry (Fig 4). These findings show that GXMR-CAR^+^ T cells can directly target *Cryptococcus* spp. in the lungs. GXMR-CAR^+^ T cells may reduce the severity of *Cryptococcus* spp. infection in the lungs (Fig. 5) through the following mechanisms: 1) production of TNF-α and IFN-γ augmenting the response of APCs and neutrophils against to *Cryptococcus* spp.; 2) release of granzyme, granulysin, and perforin by GXMR-CAR^+^ T cells, which may cause the degradation of *Cryptococcus* spp. cell walls; 3) secretion of cytokines and chemokines from GXMR-CAR^+^ T cells, which can cause epithelial cells (mucosal immunity) to produce antimicrobial peptides; and 4) fungal breakdown products taken in by dendritic cells may be cross-presented to T cells and activate humoral immunity. Our new approach to targeting *Cryptococcus* spp. with CAR^+^ T cells using an anti-GXM mAb may be extended to other invasive fungal infections using monoclonal antibodies specific to other cell wall proteins. This pioneering effort highlights the first use of CAR^+^ T cells designed to target *Cryptococcus* spp.

Cytokine release syndrome (CRS) and macrophage activating syndrome are the major limiting factors for CAR cell mediated immunotherapy. Clinicians have developed strategies to address CRS, such as treatment with Tocilizumab, an antibody to IL6R that blocks binding to the IL6 receptor, which has been successfully used in clinic. Since we are developing CAR T cells that recognize fungal antigens, at present we are not sure if there are any off target toxicities associated with this therapy. The current studies were conducted using immunosuppressive conditions by administering cyclophosphamide to eliminate the host innate immune system and induce cryptococcosis. The disadvantage of this model is that no information was collected on the CAR T-cell mediated activation of the host innate immune system for clearing infection as well as associated toxicities such as CRS. Future studies will be conducted to address these questions by developing murine homologues of GXMR-CAR and conducting studies using non-immunosuppressive conditions. The outcome of such studies will provide information about the ability of GXMR-CAR T cells to clear *Cryptococcus* infection, the participation of the host immune system in this process, and the toxicities associated with this therapy.

## Materials and Methods

### Ethics statement

All animal experiments were conducted following approval by the MD Anderson Cancer Center Institutional Animal Care and Use Committee (IACUC). Our IACUC approved protocol number is 00000555-RN01. MD Anderson’s IACUC is accredited by the Association for Assessment and Accreditation of Laboratory Animal Care (AAALAC). The IACUC was formed to meet the standards of the Public Health Service Policy on the Humane Care and Use of Laboratory Animals.

Blood samples (buffy coats) were obtained from heathy donors at the MD Anderson Blood Bank, all donors provided written informed consent. The MD Anderson Institutional Review Board (IRB) approved the protocol number LAB07-0296 for obtaining peripheral blood from healthy donors.

### Mice and *C. neoformans*

Eight-week-old female NSG mice were acquired from the Department of Experimental Radiation Oncology mouse colony at The University of Texas MD Anderson Cancer Center. The mice were maintained in a sterile biohazard facility at MD Anderson.

The *C. neoformans* yeast strain H99 (ATCC, cat. no. 208821) was allowed to grow in YPD medium until the late logarithmic phase, washed, and suspended in PBS. The concentration of yeast cells was determined using a hemocytometer, and 20 μl of a suspension of 2 × 10^6^ yeast cells/ml was used to perform intranasal inoculation of the mice.

### Construction of GXMR-CAR targeting *Cryptococcus* spp

The DNA sequence encoding the light and heavy chains of the anti-GXM mAb 18B7 clone was obtained from Casadevall et al. [26] and used to design the targeting domain of GXMR-CAR. The anti-GXM extracellular domain designed to target *Cryptococcus* spp. was fused to modified human IgG4 hinge and Fc regions [39] followed by the transmembrane and cytoplasmic domains of human CD28 and CD3-ς chains. A diagram depicting the structural regions of the GXMR-CAR construct is shown in Fig 1A. GXMR-CAR is a second-generation CAR which contains the transmembrane and intracellular regions used for CD19-specific CARs [14]. Full-length GXMR-CAR was subcloned into a lentiviral (LV) vector (Addgene; cat. no. 61422) containing the GFP sequence. The LV-GXMR-CAR construct was sequenced and verified at the Sequencing and Microarray Facility at MD Anderson.

### Generation of GXMR-CAR^+^ viral particles

To generate GXMR-CAR^+^ lentiviral particles, GXMR-CAR containing the lentiviral vectors pMD2.G (VSV-G envelope-expressing plasmid; Addgene; cat. no. 12259) and psPAX2 (second-generation lentiviral packaging plasmid; Addgene; cat. no. 12260) were transfected into HEK-293FT cells using Lipofectamine 3000 reagent (Thermo Fischer Scientific; cat. no. L3000008) according to the manufacturer’s instructions. The viral supernatant of LV-CAR– transfected HEK-293FT cells (green cells in Fig 1B) was collected every day for 3 days, and the pool of viral particles was concentrated using Lenti-X concentrator (Clontech Laboratories; cat. no. 631231). The titer of the GXMR-CAR viral particles was determined using HEK-293FT cells, and lentiviral stocks were aliquoted and maintained at −80°C.

### Transduction of cell lines and PBMCs

HEK-293FT cells (ThermoFisher; cat. no. R70007) were cultured in a six-well plate at a concentration of 5 × 10^5^ cells/ml. GXMR-CAR^+^ viral particles were added to the cells in 1 ml of RPMI 1640 medium (HyClone Laboratories) supplemented with 2 mM GlutaMAX-1 (Life Technologies; cat. no. 35050-061) and 10% heat-inactivated fetal bovine serum (HyClone Laboratories). After 24 h of incubation at 37°C, the cells were fed growth media and maintained for 2 days to evaluate the transduction efficiency via quantification of GFP expression using fluorescence microscopy and flow cytometry.

Blood samples (buffy coats) obtained from heathy donors at the MD Anderson Blood Bank were used for PBMC isolation with Ficoll-Paque PLUS solution (GE Healthcare Life Sciences; cat. no. 17144002) according to the manufacturer’s instructions. Transduction of PBMCs with LV-GXM-CAR constructs was performed using RetroNectin reagent (Takara Bio USA; cat. no. T100A/B) according to the manufacturer’s instructions. After viral transduction, human T cells were maintained in the presence of IL-2 (50 U/ml) plus IL-21 (20 ng/ml). Ten days after transduction, LV-CAR–infected T cells expressing GFP were sorted at the South Campus Flow Cytometry Facility at MD Anderson. LV-CAR T cells were expanded by stimulation with an antibody cocktail (anti-CD3/CD28; STEMCELL Technologies; cat. no. 10971) combined with IL-2 (50 U/ml) and IL-21 (20 ng/ml).

### Detection of GXMR-CAR expression on the surface of human T cells by flow cytometry

The GXMR-CAR construct expresses GFP (CAR-GFP) and a modified human IgG4 hinge region combined with an Fc region that allows for targeting by an anti-Fc antibody. The expression of GXMR-CAR on the surface of human T cells was analyzed via flow cytometry (MACSQuant; Miltenyi Biotech) using a goat anti-human IgG antibody (Fc-γ fragment-specific, 1:100 dilution; Jackson ImmunoResearch Laboratories; cat. no. 109-606-098). Human T cells were identified using flow cytometry with an anti-human CD3-PE antibody (1:100 dilution; Miltenyi Biotech; cat. no. 130-091-374). The cells double-positive for an anti-human IgG antibody and CAR-GFP represented GXMR-CAR expression on the surface of human T cells. NoDNA T cells (mock-transduced) were used as negative controls.

### Detection of interaction between GXMR-CAR^+^ T cells and the sugar moiety GXM

GXMR-CAR^+^ and NoDNA T cells were incubated with GXM (*C. neoformans* polysaccharide antigen; ZeptoMetrix) at a dilution of 1:100. An ImmunoCult human CD3/CD28 T-cell activator (STEMCELL Technologies; cat. no. 10971) was used to activate positive control T cells according to the manufacturer’s instructions. GXMR-CAR^+^ and NoDNA T cells incubated with medium alone were considered as negative controls. These cells were imaged via bright-field microscopy (Leica DMI6000B) at 400× magnification.

To determine whether GXMR-CAR^+^ or NoDNA T cells could recognize GXM, GXMR-CAR^+^ T cells that bound to GXM were identified using an anti-GXM antibody and analyzed by flow cytometry. To conduct this study, the cells were incubated for 30 min with or without *C. neoformans* polysaccharide antigens at 1:100 dilution. After washing with PBS, T cells were incubated with a murine anti-GXM mAb for 1 h (EMD Millipore; cat. no. MABF2069), washed with PBS, incubated with a goat anti-mouse IgG phycoerythrin-conjugated secondary antibody (Jackson ImmunoResearch Laboratories; cat. no. 115-116-071) and analyzed using flow cytometry. T cells without the addition of GXM served as negative controls.

### In vitro assay demonstrating that GXMR-CAR^+^ T cells target *C. neoformans* yeast

Fluorescence microscopy was used to visualize the interaction between GXMR-CAR^+^ T cells (GFP^+^) and heat killed *C. neoformans* yeast (ATCC, cat. no. 208821). The yeast were labeled with Calcofluor-white (Thermo Fischer Scientific; cat. no. L7009) according to the manufacturer’s instructions. GXMR-CAR^+^ T cells and NoDNA T cells (mock-transduced cells labeled with carboxyfluorescein succinimidyl ester) were cultured at a concentration of 1 × 10^6^ cells/ml and incubated with yeast (1 × 10^5^ cells/ml) at 37°C. After 24 h, the interaction between GXMR-CAR^+^ T cells (green) or NoDNA T cells (green) with *C. neoformans* yeast (blue) was viewed under a fluorescence microscope (Leica DMI6000B) at 100× magnification.

### Detection of *Cryptococcus* spp. in lung tissue using GMS stain and an anti-GXM immunofluorescence assay

Gomori methenamine-silver (GMS) staining and immunofluorescence assays were performed using five-micrometer lung tissue sections from mice infused with PBS, GXMR-CAR^+^ T cells, or NoDNA T cells. For immunofluorescence assay, sections were deparaffinized using a Gemini AS Automated Slide Stainer (Thermo Fisher Scientific), and antigen retrieval was performed by immersing slides in 10 mM citrate buffer (Genemed; cat. no. 10-0020) and heating for 20 min at 95°C. Tissue sections were treated with BlockAid blocking solution (Thermo Fisher Scientific; cat. no. B10710) for 30 min and incubated with an anti-GXM antibody (1:100 dilution; EMD Millipore; cat. no. MABF2069) for 1 h at room temperature. Samples were washed three times with PBS, incubated with anti-mouse IgG1-fluorescein isothiocyanate for 45 min at room temperature (1:100 dilution; BD Pharmingen; cat. no. 553443), and washed three times with PBS. ProLong Gold Antifade Mountant (Invitrogen; cat. no. 36935) was added to the tissue prior to sealing with a coverslip, and imaging of lung tissue was performed at 400x using the Vectra^®^ Polaris^™^ Automated Quantitative Pathology Imaging System (Perkin Elmer) located at the MD Anderson Flow Cytometry and Imaging Core Facility in Houston, TX. The inForm^®^ Tissue Finder^™^ and inForm^®^ Cell Analysis^™^ software (Perkin Elmer) was utilized to quantitate the region of GXM positive staining and the total area of lung tissue on each slide. To conduct the inForm analysis, the tissue was segmented using a large pattern scale, fine segmentation resolution, edges were trimmed by 5 pixels, and a minimum segment size of 1550, and the components for the training set were DAPI, autofluorescence, and opal 520 for GXM-FITC. The tissue categories were: GXM region (red), lung tissue (green), non-tissue (blue). The percentages of the “area of GXM positive staining / total area of lung tissue” of each mouse per group were compared using ANOVA. Pair-wised comparisons were conducted using Tukey’s HSD (honestly significant difference) test. Statistical significance was defined as P value < 0.05. All analyses were performed using R version 3.5.1 [40]. To conduct an ImageJ analysis of the GXM positive cells in the lungs detected by immunofluorescence, the images were acquired at 200× magnification using an Olympus CKX41 microscope. The color threshold mode of ImageJ software was used for detection of fluorescent GXM stained cells and the threshold was adjusted to include areas of GXM positive cells. The analyze particles mode was applied to remove background, and the area of Titan cells was obtained for each image. The results are expressed as pixels^2^ and the same area of the tissue was analyzed for each group.

## Acknowledgments

We thank Dr. Jared Burks, Co-Director of the MD Anderson Flow Cytometry and Imaging Core Facility supported by NIH/NCI under award number P30CA016672 for conducting the imaging of GXM positive lung tissue and performing automated analysis of the tissue using the inForm Tissue Finder software. TAdS was financially supported by the São Paulo Research Foundation, Brazil (grant numbers 2016/23044-1 and 2016/04877-2). Funding for PK and PJH was provided by the Department of Pediatrics Research, MD Anderson and by the National Institute of Allergy and Infectious Disease (R21 grant AI127381-02). The funders had no role in study design, data collection and analysis, decision to publish, or preparation of the manuscript.

## Author Contributions

1. Conceptualization: KP, TAdS
2. Data curation: KP, TAdS, PJH
3. Formal analysis: KP, TAdS, PJH, QW
4. Funding acquisition: KP, TAdS,
5. Investigation: KP, TAdS, PJH
6. Methodology: KP, TAdS, PJH
7. Project administration: KP, TAdS, PJH
8. Resources: KP, TAdS
9. Software: KP, TAdS, PJH, QW
10. Supervision: KP
11. Validation: KP, TAdS, PJH
12. Visualization: KP, TAdS, PJH
13. Writing – original draft: KP, TAdS, PJH
14. Writing – review & editing: KP, TAdS, PJH, QW, IB, TL, AM

---

**Fig S1.**
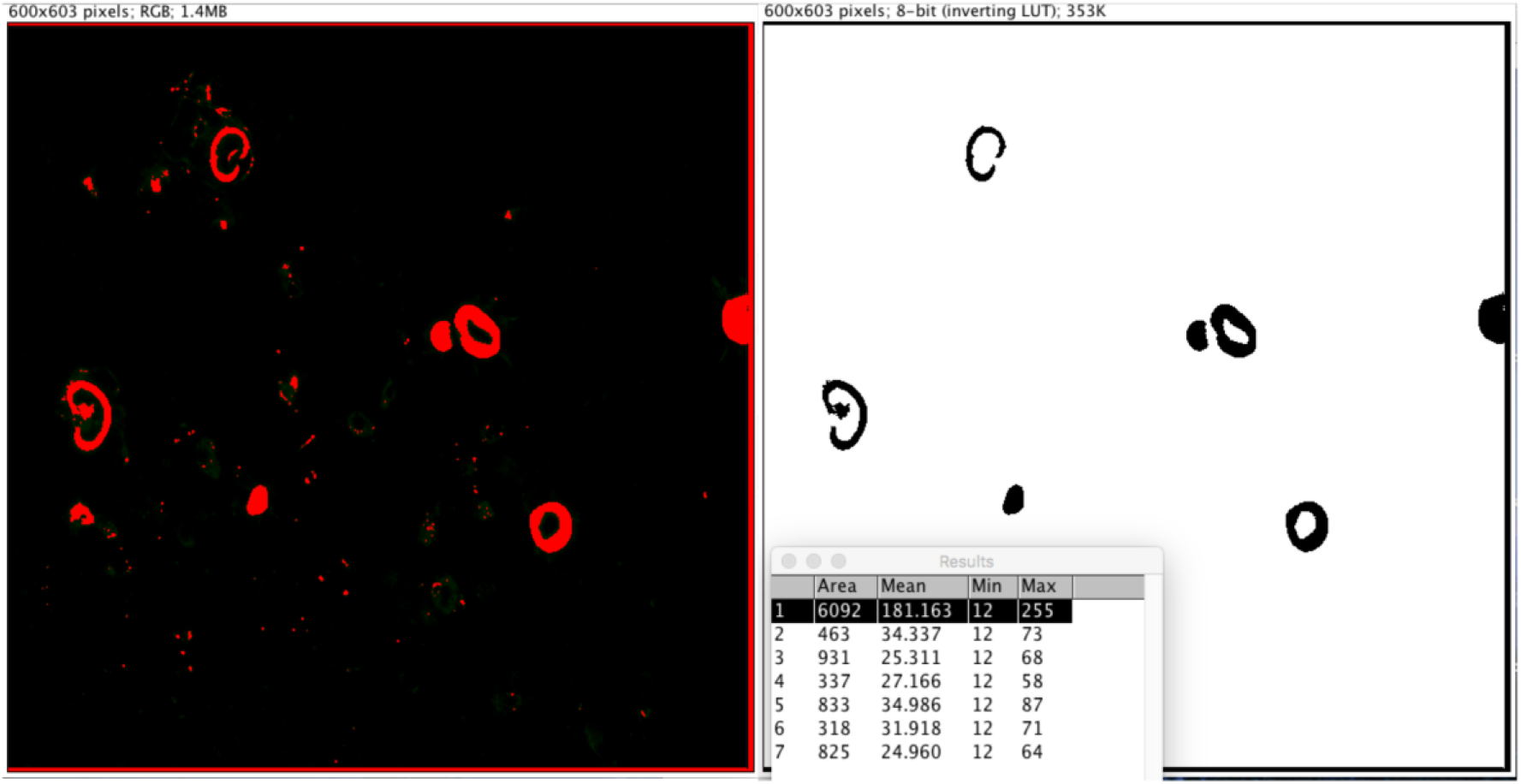
Processing steps used to quantify area of GXM stained Titan cells in anti-GXM immunofluorescence assay by using ImageJ. ImageJ software was used to quantify the area of GXM positive staining in lungs from the nine the images that are shown in Fig 5A.

